# Neural representations of social valence bias economic interpersonal choices

**DOI:** 10.1101/355115

**Authors:** Paloma Díaz-Gutiérrez, Juan E. Arco, Sonia Alguacil, Carlos González-García, María Ruz

## Abstract

Prior personal information is highly relevant during social interactions. Such knowledge aids in the prediction of others, and it affects choices even when it is unrelated to actual behaviour. In this investigation, we aimed to study the neural representation of positive and negative personal expectations, how these impact subsequent choices, and the effect of mismatches between expectations and encountered behaviour. We employed functional Magnetic Resonance Imaging in combination with a version of the Ultimatum Game (UG) where participants were provided with information about their partners’ moral traits previous to their fair or unfair offers. Univariate and multivariate analyses revealed the implication of the supplementary motor area (SMA) and inferior frontal gyrus (IFG) in the representation of expectations about the partners in the game. Further, these regions also represented the valence of expectations, together with the ventromedial prefrontal cortex (vmPFC). Importantly, the performance of multivariate classifiers in these clusters correlated with a behavioural choice bias to accept more offers following positive descriptions, highlighting the impact of the valence on the expectations on participants’ economic decisions. Altogether, our results suggest that expectations based on social information guide future interpersonal decisions and that the neural representation of such expectations in the vmPFC is related to their influence on behaviour.

## 1. Introduction

Decision-making is a crucial constituent of our daily life. To make choices that best fit our goals, we must rapidly weight different sources of information in an efficient manner. An elegant approach to understand how we perform such weighting comes from the framework of predictive coding (Friston, 2005), where optimal decision-making combines sensory input (*evidence*) with predictions (*priors;* Schwarz et al., 2016; Summerfield and De Lange, 2014). The role of these predictions has been thoroughly examined in non-social decisions, where several studies have shown pre-activation of target-related brain areas during the expectation period, prior to target onset (e.g., Esterman and Yantis, 2010; González-García et al., 2016; Puri et al., 2009). However, a large part of decisions involve social contexts, where we constantly engage in interactions with others. Still, the role of expectations in such scenarios remains unclear.

When making decisions in complex scenarios, people tend to choose more often and faster the options that match their personal preferences (with higher personal value) even when the objective task value of the different alternatives is similar (Lopez-Persem et al., 2016). This leads to suboptimal decisions that do not properly consider potential future outcomes (Fleming, Thomas, & Dolan, 2010). This is also the case for interpersonal decisions, which can be biased by several sources of information at different stages of processing (Díaz-Gutiérrez, Alguacil, & Ruz, 2017). For instance, in the Ultimatum Game (UG; Güth, Schmittberger, & Schwarze, 1982; Moser, Gaertig, & Ruz, 2014), participants receive monetary offers from game partners and decide whether to accept them or not. Acceptance leads to both parts earning their split; whereas no gains are earned after a rejection. Here, “rational” decisions from an economic point of view should be of acceptance, since you can only earn money. However, choices are strongly influenced by the fairness of the offer (how balanced both halves of the split are). People often show high rejection rates towards unfair offers (Sanfey, Rilling, Aronson, Nystrom, & Cohen, 2003), which has been explained in terms of inequity-aversion tendencies (Fehr & Camerer, 2007) and punishment (Brañas-Garza, Espín, Exadaktylos, & Herrmann, 2014). Others have emphasized the importance of social norms, and how these impact the perception of fairness (Chang & Sanfey, 2013). In these scenarios, the mechanisms underlying the processing of offers depending on their fairness and participants’ subsequent responses have been extensively studied. Here the role of the anterior cingulate cortex (ACC) and supplementary motor area (SMA) stands out, concerning both fairness and people’s decisions (for a meta-analysis, see Gabay et al., 2014). Authors such as Sanfey et al., (2003) have shown the involvement of the anterior insula (aI) in fairness processing. Also, Corradi-Dell’Acqua, Civai, Rumiati, & Fink (2013) differentiated its role from the one of the medial prefrontal cortex (mPFC), which appears to be linked to emotional self-related responses during interpersonal bargaining situations.

Despite the extensive and diverse studies in interpersonal games, it is largely unknown how the brain represents socially relevant priors in these scenarios. Recent proposals have tried to link predictive coding and the representation of social traits in relation to social expectations (e.g., Tamir and Thornton, 2018). Several studies have described a set of regions underlying the representation of knowledge that guides social predictions in a broad context (termed Social Cognition Network; Frith & Frith, 2008), including personal traits, stereotyping, semantic knowledge about people or inferences about others and their mental states (Tamir & Thornton, 2018; Tamir, Thornton, Contreras, & Mitchell, 2016). This network includes the temporoparietal junction (TPJ), superior temporal sulcus (STS), precuneus (PC), anterior temporal lobes (ATL), amygdala and the mPFC (Contreras et al., 2013; Frith, 2007; Frith and Frith, 2001; Mitchell et al., 2008). These regions underlie processes such as Theory of Mind (ToM; Saxe and Kanwisher, 2003). Similarly, in decisions in social contexts, the mPFC has been related to expectations about others’ behaviour (Corradi-Dell’Acqua, Turri, Kaufmann, Clément, & Schwartz, 2015). Importantly, prior expectations during social decisions also influence behaviour when they are not followed by their usual consequences. In this line, different studies (Fouragnan et al., 2013; Ruz and Tudela, 2011) have observed increased activation in brain areas associated with cognitive control, such as the ACC and the aI when expectations about partners do not match their subsequent behaviour. Similarly, Chang and Sanfey (2013) found a relationship between the deviation of the expectations and increased activation in the aI, ACC and SMA. Specifically, in the UG, an increase of activation in the dorsolateral PFC (dlPFC) and aI has been related to participants’ reaction to unfair offers (Knoch, Pascual-Leone, Meyer, Treyer, & Fehr, 2006; Sanfey et al., 2003), which has also been interpreted as a violation of what we expect from others.

In addition to this, social expectations can also be based on the personal traits of others, which are an essential component of social representations (Tamir & Thornton, 2018). The priors that they generate relate to stereotypes and interact with perceptual processes (Stolier & Freeman, 2016, 2017). These personality traits can be decomposed in three different dimensions: rationality, social impact and, crucially to our investigation, valence (positive vs. negative; Tamir and Thornton, 2018; Thornton and Mitchell, 2017). The representation of the character of others in association with positive or negative information is an important source of bias in interpersonal decisions (Díaz-Gutiérrez et al., 2017). For instance, Delgado et al. (2005), found that participants trusted partners associated with positive moral traits more than those having negative ones. Furthermore, a variety of studies employing the UG paradigm have observed that participants tend to accept more offers from partners associated with positive descriptions, compared to negative ones (Gaertig, Moser, Alguacil, & Ruz, 2012). This tendency is steeper when participants navigate uncertain scenarios (Ruz, Moser, & Webster, 2011). Moreover, in this context, the use of high-density electroencephalography (EEG) has shown that negative descriptions of partners lead to a higher amplitude of the medial frontal negativity (MFN; associated with the evaluation of outcomes, Hajcak et al., 2006; Yeung and Sanfey, 2004) when decisions are made (Moser et al., 2014). These data indicate how, regardless of fairness, people evaluate offers as more negative when they come from a disagreeable partner. Such knowledge about personal traits has been suggested to be integrated by the mPFC (Van Overwalle, 2009). For example, this area increases its coupling with other regions responding to specific traits (Hassabis et al., 2014), and shows heightened activation when a partner’s behaviour violates previous trait implications (Ma et al., 2012).

Nonetheless, despite the key relevance of valence in psychological theories and its marked impact on social decision-making, it is not well understood how valence is represented at the neural level and its effect on subsequent choices (Barrett & Bliss-Moreau, 2009). Results of a recent meta-analysis (Lindquist, Satpute, Wager, Weber, & Barrett, 2015) provide evidence of a general recruitment of a set of regions for valenced versus neutral information, including the bilateral aI, the ventral and dorsal portions of the mPFC (vm/dmPFC), the dorsal ACC, SMA, and lateral PFC. Lindquist et al. (2015) found that the vmPFC/ACC was more frequently activated in positive vs. negative than in positive vs. neutral contrasts, which could indicate that these regions represent valence information along a single bipolar dimension.

Taking all this into account, in the current functional Magnetic Resonance Imaging (fMRI) study, we employed a modified version of the UG (Gaertig et al., 2012) to investigate how socially relevant priors represented by the valence of personal descriptions of partners bias interpersonal economic choices. First, we aimed to study which neural regions code for the generation and maintenance of positive and negative expectations about other people. In a second step, we assessed how these expectations bias decisions. We expected to find specific neural representations underlying the expectations about the partners, with different patterns depending on the valence of these predictions (Lindquist et al., 2015). Specifically, we hypothesized that these patterns would be represented in regions related to social cognition and priors in decision-making (Contreras et al., 2012; González-García et al., 2016; Saxe & Kanwisher, 2003). Last, we intended to ascertain which neural mechanisms were engaged when there is a mismatch between personal expectations and the partners’ behaviour. We predicted that control-related areas would be engaged when the valenced description was not congruent with the subsequent partner’s behaviour.

## 2. Methods

### 2.1. Participants

Twenty-four volunteers were recruited from the University of Granada (M = 21.08, SD = 2.92, 12 men), matching the sample size employed in Moser et al. (2014), who implemented the same version of the task for electroencephalography (EEG). This sample is similar to previous fMRI studies using the UG (Chang and Sanfey, 2013; Grecucci, Giorgetta, Bonini & Sanfey, 2013). All participants were right-handed with normal or corrected vision and received economic remuneration (20-25 Euros, proportionally to their acceptance rates). Participants signed a consent form approved by the Ethics Committee of the University of Granada.

### 2.2. Apparatus and stimuli

We employed 16 adjectives used in previous studies (Gaertig et al., 2012; Moser et al., 2014; Ruz et al., 2011; see Table 1) as trait-valenced descriptions of the game proposers, extracted from the Spanish translation of the Affective Norms for English Words database (ANEW; Redondo et al., 2007). Half of the adjectives were positive (M = 7.65 valence, SD = 0.43), and the other half were negative (M = 2.3 valence, SD = 0.67). All words were matched in arousal (M = 5.69, SD = 0.76), number of letters (M = 6.19, SD = 1.42) and frequency of use (M = 20.19, SD = 18.47). In addition, we employed numbers from 1 to 9 (two in each trial) in black colour to represent different monetary offers. Stimuli were controlled and presented by E-Prime software (Schneider, Eschman, & Zuccolotto, 2002). Inside the scanner, the task was projected on a screen visible to participants through a set of mirrors placed on the radiofrequency coil.

**Table 1.** List of adjectives employed in the task (Gaertig et al., 2012).

| Positive words | Negative words |
| --- | --- |
| Friend | Criminal |
| Generous | Cruel |
| Honest | Disloyal |
| Honourable | False |
| Humble | Guilty |
| Kind | Hostile |
| Loyal | Selfish |
| Warm | Traitor |

### 2.3. Task and procedure

To add credibility to the interpersonal game setting, participants were told that they were about to receive offers made by real participants in a study of a previous collaboration with a foreign university. Furthermore, to engage participants in the game as a real social scenario, prior to the scanner they performed two tasks in which they had to make economic offers that would be used for other participants in future studies. In one of the tasks, participants acted as proposers, filling a questionnaire where they had to make offers for 16 different unknown partners, who would be involved in future experimental games. Here, they had to split 10 Euros into two parts, one for themselves and the other for their partners. Additionally, in a second task, they played a short version of the Dictator Game (Kahneman, Knetsch, & Thaler, 1986), where they decided how to divide another 10 Euros between themselves and an anonymous partner, who would have a merely passive role concerning the output of the offer. Moreover, participants were told that the offers that they were about to see in the scanner were each provided by a different partner who previously performed the same tasks as they did before the scanner, and therefore, the offers were real examples of other participants’ responses when acting as proposers. Participants were informed that each offer would be preceded by a word that had been obtained as an output from a series of personality and social questionnaires filled by their partners and, therefore, that these adjectives described them in some way (see Table 1). Choices made by participants had an influence in their final payment, as it actually varied (20-25 Euros) according to their choices during the game in the scanner. In a post-scanning informal debriefing session, none of the participants reported suspicions regarding the background story of this procedure, which has also been used successfully in other settings (e.g. Correa, Alguacil, Ciria, Jiménez, & Ruz, 2020; Correa et al., 2017).

In the scanner, participants played the role of the responder in a modified UG (e.g., Gaertig et al., 2012), deciding whether to accept or reject monetary offers made by different partners (proposers). If they accepted the offer, both parts earned their respective splits, whereas if they rejected it, neither of them earned money from that exchange. Offers consisted of splits of 10 Euros, which could be fair (5/5, 4/6) or unfair (3/7, 2/8, 1/9). The number presented at the left on the screen was always the amount of money given to the participant, and the one on the right side was the one proposed by the partners for themselves.

Personal information about the partners was included as adjectives with different valence. A third of these descriptions was positive, another third negative, and the last third was neutral, represented by text indicating the absence of information about that partner (“no test”). The valence of the adjectives was orthogonal to the fair-unfair nature of the offer. The order of the offers and adjectives was randomized, and each type of personal information (positive, negative, no information) preceded each offer equally within and across runs. Decision-response associations were counterbalanced between participants.

Participants performed a total of 192 trials, arranged in 8 runs (24 trials per run). In each run, a start cue of 6 s was followed by 24 trials. Each trial (see Figure 1) started with an adjective for 1 s (mean = 2.98º), preceding a jittered interval lasting 5.5 s on average (4-7 s, +/0.76º). Then, the offer appeared for 0.5 s (1.87º), followed by a second jittered interval (mean = 5.5 s; 4-7 s, +/0.76º). Overall, each run lasted 5.1 minutes and the whole task 41 minutes approximately.

**Figure 1.**
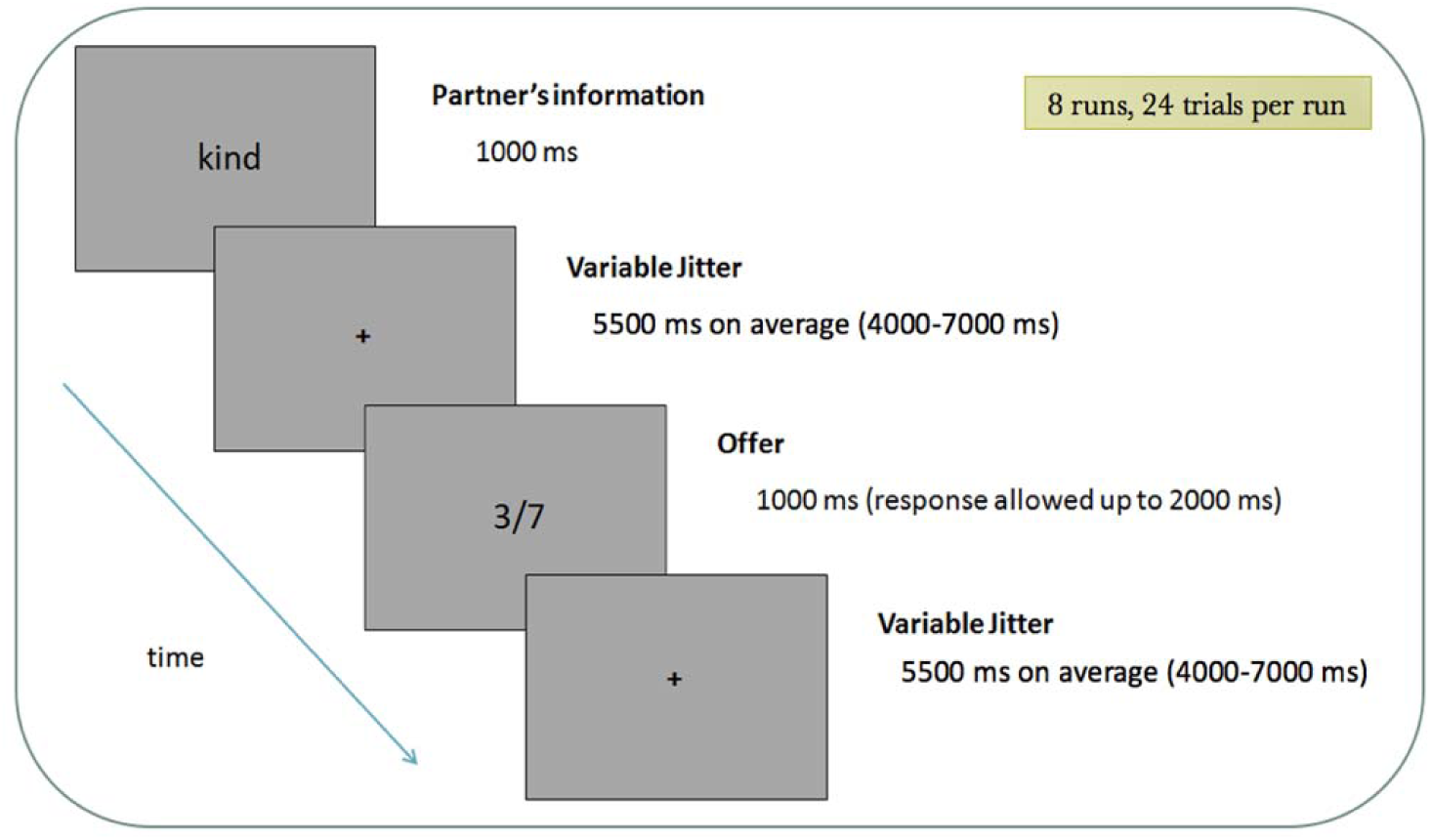
Sequence of events in a trial. The task varied the Valence of the partner’s information (Positive, Negative, No information) and the Fairness of the offer (Fair/Unfair), which were manipulated orthogonally in the design.

### 2.4. Image acquisition and preprocessing

MRI images were acquired using a Siemens Magnetom TrioTim 3T scanner, located at the Mind, Brain and Behavior Research Center in Granada. Functional images were obtained with a T2*-weighted echo-planar imaging (EPI) sequence, with a TR of 2000 ms. Thirty-two descendent slices with a thickness of 3.5 mm (20% gap) were extracted (TE = 30 ms, flip angle = 80 º, voxel size of 3.5 mm^3^). The sequence was divided into 8 runs, consisting of 166 volumes each. After the functional sessions, a structural image of each participant with a high-resolution T1-weighted sequence (TR = 1900 ms; TE = 2.52 ms; flip angle = 9º, voxel size of 1 mm^3^) was acquired.

Data were preprocessed with SPM12 software (http://www.fil.ion.ucl.ac.uk/spm/software/spm12/). The first three volumes of each run were discarded to allow the signal to stabilize. Images were realigned and unwarped to correct for head motion, followed by slice-timing correction. Afterwards, T1 images were coregistered with the realigned functional images. Then, functional images were spatially normalized according to the standard Montreal Neurological Institute (MNI) template and smoothed employing an 8 mm Gaussian kernel. Low-frequency artefacts were removed using a 128 high-pass filter. Data for multivariate analyses was only head-motion and slice-time corrected and coregistered.

### 2.5. Univariate analyses

First-level analyses were conducted for each participant, following a General Linear Model in SPM12. We employed an event-related design, where activity was modelled using regressors for each valence type of adjective and for the offers. The estimated model included three regressors for the Words (positive, negative, no information) and six for the Offers (Fair offers_Positive, Fair offers_Negative, Fair offers_Neutral, Unfair offers_Positive, Unfair offers_Negative, Unfair offers_Neutral). Note that since decisions were made when the offers appeared, and that responses (choices) showed a strong dependency on offer fairness, offer fairness and decisions cannot be modelled separately. Given our research questions, we modelled the offer events considering their fairness regardless of participants’ choices. Regressors were convolved with a standard hemodynamic response, with adjectives modelled with their duration (1 s + jitter), and offers modelled as events with zero duration. This temporal difference is accounted by the fact that the words describing the partners trigger preparatory processes, which extend in time (e.g. Bode & Haynes, 2009; Di Russo et al., 2017; González-García, Arco, Palenciano, Ramírez, & Ruz, 2017; González-García et al., 2016; Sakai, 2008), whereas the processing of the offers ends shortly after with the response of each trial (see Moser et al., 2014). In addition, the orthogonal manipulation of these variables in the design avoided covariance confounds between word cues and target offers.

At the second level of analysis, *t*-tests were conducted for comparisons related to the presence of expectations (information about the partner > no information), the valence of the information (positive > negative, negative > positive) and the fairness of the offer (fair > unfair, unfair > fair). We also carried out contrasts for congruence effects between the events, where we had congruent (positive descriptions followed by fair offers, negative descriptions followed by unfair offers) and incongruent trials (positive descriptions followed by unfair offers, negative descriptions followed by fair offers). To control for false positives at the group level, we employed permutations tests with statistical non-parametric mapping (SnPM13, http://warwick.ac.uk/snpm) and 5000 permutations. We performed cluster-wise inference on the resulting voxels with a cluster-forming threshold of 0.001, which was later used to obtain significant clusters (FWE corrected at *p<*0.05).

### 2.6 Multivariate analyses

We performed MVPA to examine the brain areas representing the valence of the expectations, that is, the regions containing information about whether the partners were described with positive vs. negative adjectives. To this end, we performed a whole-brain searchlight (Kriegeskorte et al., 2006) on the realigned images (prior to normalization). We employed The Decoding Toolbox (TDT; Hebart et al., 2015), to create 12-mm radius spheres, where linear support vector machine classifiers (C=1; Pereira et al., 2009) were trained and tested using a leave-one-out cross-validation scheme, employing the data from the 8 scanning runs (training was performed with data from 7 runs and tested in the remaining run, in an iterative fashion). We used a Least-Squares Separate model (LSS; Turner, 2010) to reduce collinearity between regressors (Abdulrahman & Henson, 2016; Arco et al., 2018). This approach fits the standard hemodynamic response to two regressors: one for the current event of a trial (positive/negative adjective) and a second one for all the remaining events and trials. As in the previous analyses, adjective regressors were modelled with their duration (1 s + jitter) and offers with zero duration. Consequently, the output of this model was one beta image per event (total = 128 images, 64 for each type of adjective, 112 for training and 16 for testing in each iteration). Afterwards, at the group level, non-parametrical statistical analyses were performed on the resulting accuracy maps following the method proposed by Stelzer et al. (2013) for MVPA data. We permuted the labels and trained the classifier 100 times for each participant. The resulting maps were then normalized to an MNI space. Afterwards, we randomly picked one of these maps per each participant and averaged them, obtaining a map of group accuracies. This procedure was repeated 50000 times, building an empirical chance distribution for each voxel position and selecting the 50th greatest value, which corresponds to the threshold that marks the statistical significance. Only the voxels that surpassed this were considered significant. The resulting map was FWE corrected at 0.05, computing previously the cluster size that matched this value from the clusters obtained in the empirical distribution.

Importantly, the valence of the description influenced acceptance rates, which could generate potential confounds in the previous decoding. The association between hand and decision (left/right, acceptance/rejection) was fully counterbalanced across participants, but remained constant for each of them. Therefore, the classifier could use response information (accept vs. reject) when decoding valence. To clarify this issue, we performed a response classification at the offer period (following the same procedure as for the valence decoding). Then, we ran a conjunction analysis, computing the intersection between valence and response group maps to examine whether the regions containing relevant information about the valence were the same as those representing the participants’ decisions (accept vs. reject). Moreover, to test additionally the potential overlap between the neural representations of participants’ decisions and the valence of the expectations about the partners, we performed a cross-classification analysis (Kaplan, Man, & Greening, 2015) between these two domains. Following again the same classification procedure described above in this section, we trained the classifier with the participants’ responses to the offers (accept vs. reject) and tested it on the valence of the partner’s descriptions (positive vs. negative).

### 2.7. Relationship between decoding accuracy and choices

To examine the extent to which the fidelity of representation of (positive vs. negative) personal priors relates to the decisions made by participants, we performed a correlation analysis between an individual bias index and mean decoding accuracy values from each significant cluster in the MVPA described above. To obtain this behavioural index, for each participant we subtracted the average acceptance rate following negative descriptions from the average acceptance rate after positive descriptions (regardless of the nature of the offer). For each subject, we performed a one-tailed (right) Spearman’s correlation between the behavioural index and the decoding accuracy from each significant cluster (Bonferroni-corrected for multiple comparisons). To further ascertain that participants’ motor responses were not contaminating this link between valence representation and interpersonal choices, we ran an additional correlation analysis following the same approach, this time to examine the link between valence’ decoding results and the response made by participants (acceptance or rejection of the offer). Therefore, for each participant, we calculated their average acceptance rate in general, regardless of the valence of the expectation and the fairness of the offers.

## 3. Results

### 3.1. Behavioural data

Acceptance rates (AR) and reaction times (RTs) were analysed in a Repeated Measures ANOVA, with Offers (fair/unfair) and Valence of the descriptions (positive, negative, neutral) as factors. The Greenhouse-Geisser correction was applied whenever the sphericity assumption was violated.

#### 3.1.1. Acceptance rates

Participants responded on 100% of the trials. Data showed (see Figure 2) a main effect of Offer *F*_1,23_ = 74.50, *p* < .001, η_p_^2^ = .764, where fair offers were accepted more often (*M* = 84.09%; *SD* = 22.10) than unfair ones (*M* = 24.18%; *SD* = 24.10). Valence was also significant, *F*_2,22_ = 13.735, *p =* .001, η_p_^2^ = .374. Participants accepted more offers when they were preceded by a positive description of the partner (*M* = 59.39%; *SD* = 23.09), than when there was no information (*M* = 56.31%; *SD* = 21.89) or when this was negative (*M* = 46.70%; *SD* = 24.33). Planned comparisons revealed that these differences were significant between all pairs (all *p*_s_<.05). Finally, the Offer X Valence interaction was also significant, *F*_2,22_ = 4.262, *p =* .033, η_p_^2^ = .156. Planned comparisons showed that for fair offers, there were differences between all comparisons (*ps* = .002) except between positive and neutral information (*p =* .399), whereas for unfair offers, there was no difference in acceptance rates between negative and neutral information (*p* = .074) but there was for the rest of the pairwise comparisons: *ps* < .01)

**Figure 2.**
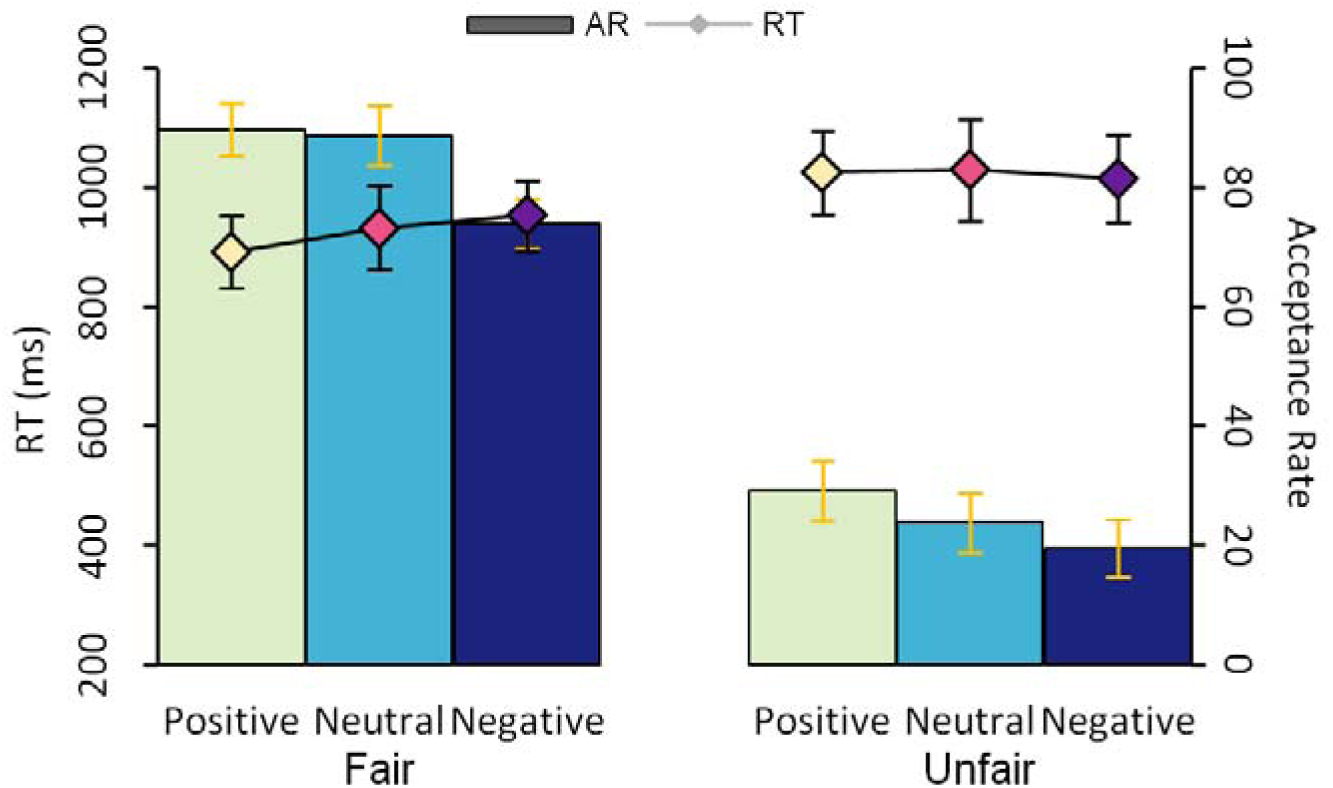
Acceptance Rates (AR, bars) and reaction times (RT, lines) to fair and unfair offers preceded by positive, negative and neutral descriptions of the partner (error bars represent S.E.M).

#### 3.2.1. Reaction times

Results showed (see Figure 2) a main effect of Offer *F*_1,23_ = 22.489, *p* < .001, η_p_^2^ = .494, where participants took longer to respond to unfair (*M* = 1023.53 ms; *SD* = 373.10 ms) than to fair offers (*M* = 925.62 ms; *SD* = 309.57 ms). Neither Valence, *F*_2,22_ = 1.05, *p =* .341, or its interaction with Fairness, *F*_2,22_ = 1.956, *p =* .168 were significant. In addition, to measure the influence of expectations on participant’s responses (see Ruz et al., 2011), we ran an ANOVA where we included the valence of the descriptions and the decision (accept, reject) made to the offers. Here, we did not find any effect of Valence, *F*<1, but we found significant effects of Decision, *F*_1,23_ = 5.519, *p =* .028, η_p_^2^ = .194, since participants were faster to accept (*M* = 951.37 ms; *SD* = 356.01 ms) than to reject the offers (*M* = 988.97 ms; *SD* = 316.91 ms). Interestingly, data showed an interaction Valence X Decision, *F*_2,22_ = 4.23, *p =* .025, η_p_^2^ = .155, replicating previous findings (Gaertig et al., 2012; Ruz et al., 2011). Planned comparisons indicated that these differences in RT for responses took place only after positive, *F*_1,23_ = 13.997, *p =* .001, η_p_^2^ = .378 (Accept: *M* = 927.60 ms, *SD* = 297.37 ms; Reject: *M* = 993.91 ms, *SD* = 335.52 ms), and neutral descriptions, *F*_1,23_ = 4.504, *p =* .045, η_p_^2^ = .165 (Accept: *M* = 955.8 ms, *SD* = 304.96 ms; Reject: *M* = 987.80 ms, *SD* = 328.48 ms), but not for negative descriptions, *F*<1.

### 3.2. Neuroimaging data

#### 3.2.1. Univariate results

##### Expectations

During the presentation of the description and the time interval that followed, that is, when participants had personal information to **generate expectations** [(Positive adjective & Negative adjective) > No Information], we observed a cluster of activity (see Figure 3a) in the left dorsal aI (*k =* 109; -33, 21, 4) and bilateral Supplementary Motor Cortex (SMA; *k =* 138; -8, 11, 53; see Fig. 3). Additionally, the right inferior parietal lobe (right IPL) showed higher activity (*k =* 264; 55, -35, 53) for **positive descriptions** compared to negative ones. No cluster surpassed the statistical threshold (*p*>0.05) for the opposite contrast.

**Figure 3.**
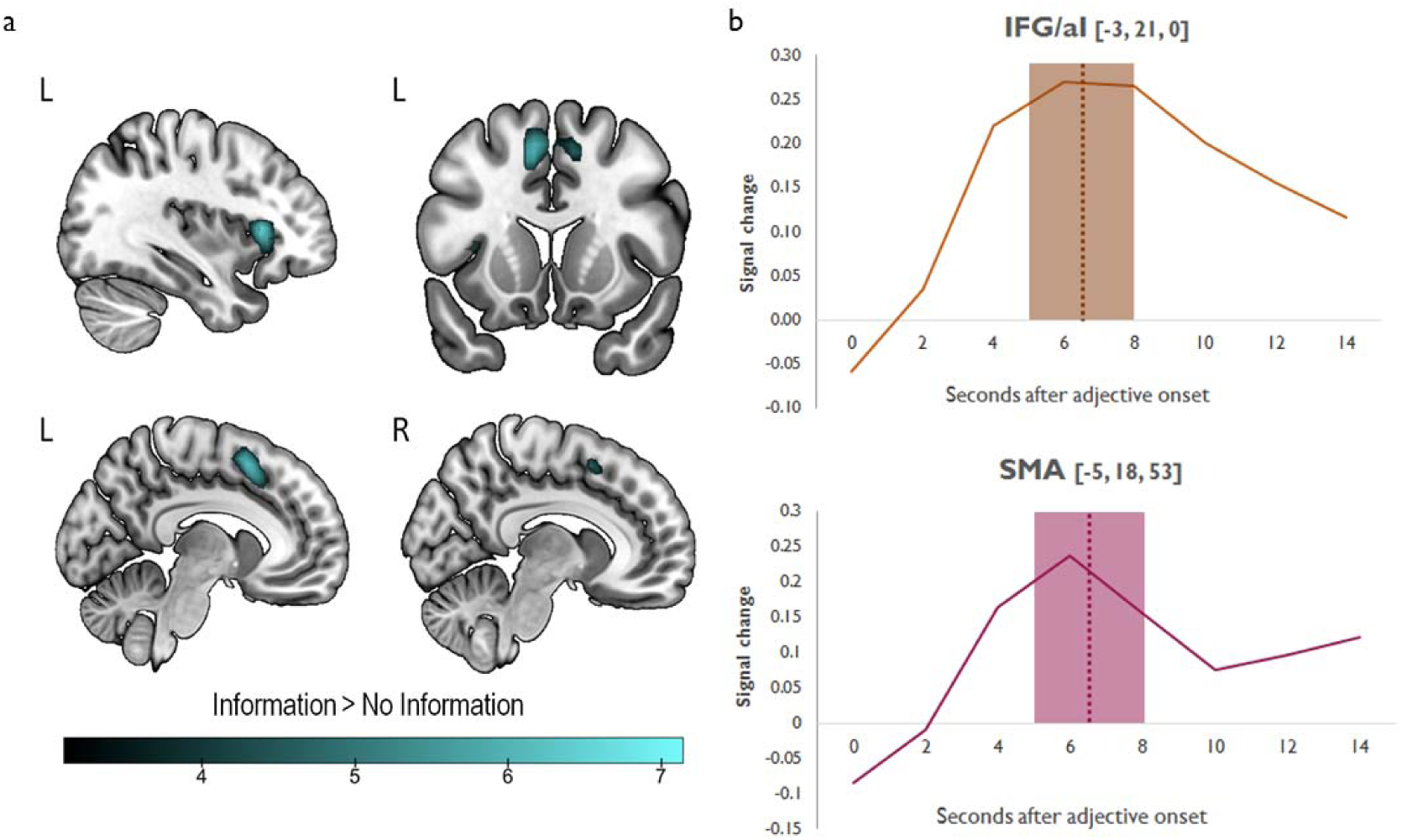
**a)** Univariate results during the expectation period. Scales reflect peaks of significant t-values (p<.05, FWE-corrected for multiple comparisons). **b)** Time course of activation in the IFG/aI (−3, 21, 0; *top*) and SMA (−5, 18, 53; *bottom*) clusters obtained from the conjunction analysis. From these regions, we extracted the signal change values related to the processing of personal information minus the average during the neutral condition, time-locked to the adjective onset. The shaded areas show the variable time window during which the offer could appear (5-8 s after the adjective onset) whereas the dotted lines show its average (6.5 after the adjective onset).

During *offer processing*, the previous presentation of **personal information** about the partner [(Offer_Pos & Offer_Neg > Offer_Neu] yielded again significant activity involving the bilateral dorsal aI and right SMA (*k =* 23349; -33, 21, 4).

To check whether the regions related to personal information were the same during the presentation of the valenced adjectives and during the presentation of the offer (positive and negative > neutral in both cases), we ran a conjunction analysis with the regions significant in both contrasts (Nichols, Brett, Andersson, Wager, & Poline, 2005). Similar to each contrast individually, we observed two clusters: one in the left IFG/aI (*k* = 93; -3, 21, 0) and one involving bilateral SMA (*k* = 126; -5, 18, 53), suggesting that both areas increased their activation during the expectation and offer stages (see Figure 3b).

##### Offer fairness

**Fair offers** (Fair > Unfair) generated activity (see Figure 4) in the right medial frontal gyrus (mFG) and ACC (*k =* 171; 6, 39, -14), while the opposite contrast (unfair > fair) did not yield any significant clusters (*p*>0.05). Furthermore, we examined neural responses depending on whether previous expectations were matched or not by the nature (fair vs. unfair) of the offer. Here, **congruence** (see Figure 4) between expectations and offer (Congruent > Neutral) showed a cluster of activity in right cerebellum (right Crus; *k* = 153; 17, -88, -32). Conversely, **incongruence** (see Figure 4) between expectations and offer (Incongruent > Neutral) yielded activations in the right medial Superior Frontal Gyrus (mSFG) and its lateral portion bilaterally (*k* = 401; 13, 39, 56), as well as in left IFG (*k* = 177; -54, 39, 0). Lastly, regarding general conflict effects, a comparison between **mismatch** (incongruent) **vs. match** (congruent) trials showed clusters of bilateral activity in the IFG/aI (*k =* 232; -43, 25, -11/ *k =* 140; 34, 35, 4; see Figure 4).

**Figure 4.**
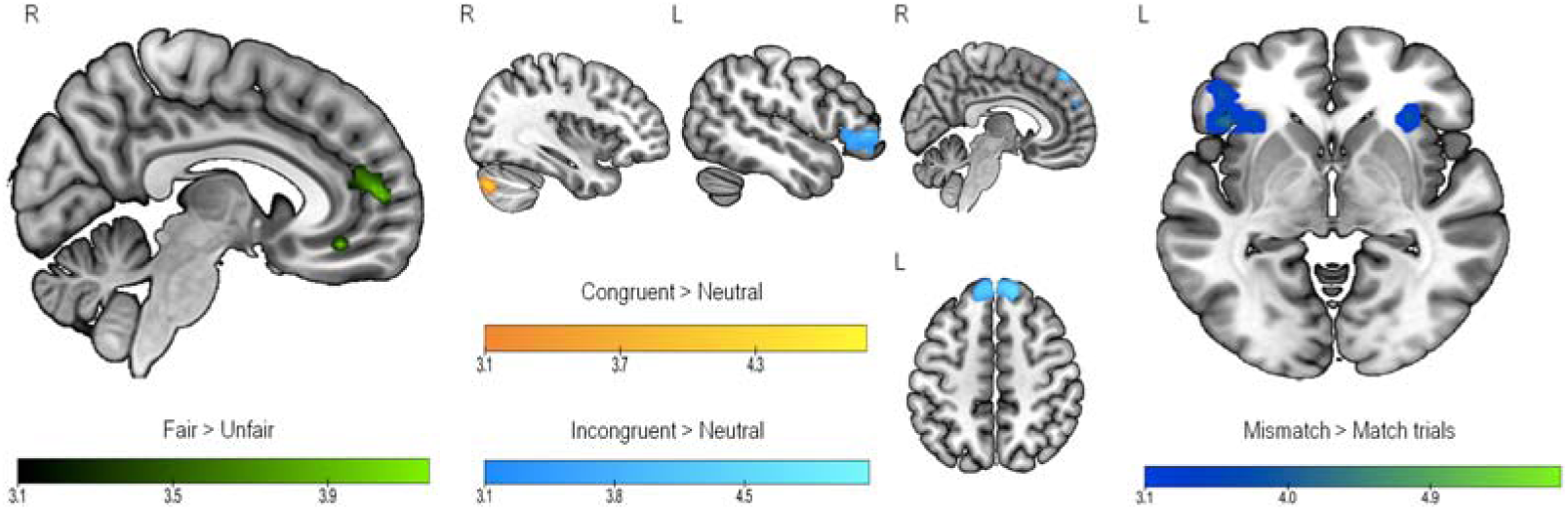
Univariate results for the offer. Scales reflect peaks of significant t-values (p<.05, FWE-corrected for multiple comparisons).

#### 3.2.2. Multivariate results

##### Valence of expectations’ classification

Expectations about the partners (positive vs. negative information) showed distinct patterns of neural activity in a cluster including the left inferior and middle frontal gyrus (IFG/MFG) and aI (*k* = 319; -46.5, 28, -32.2), the bilateral ventromedial prefrontal cortex (vmPFC) and ACC (*k* = 483; 6, 21, -19.6), and the bilateral middle cingulate cortex (MCC) and SMA (*k =* 339; -4.5, 14, 35; see Figure 5).

**Figure 5.**
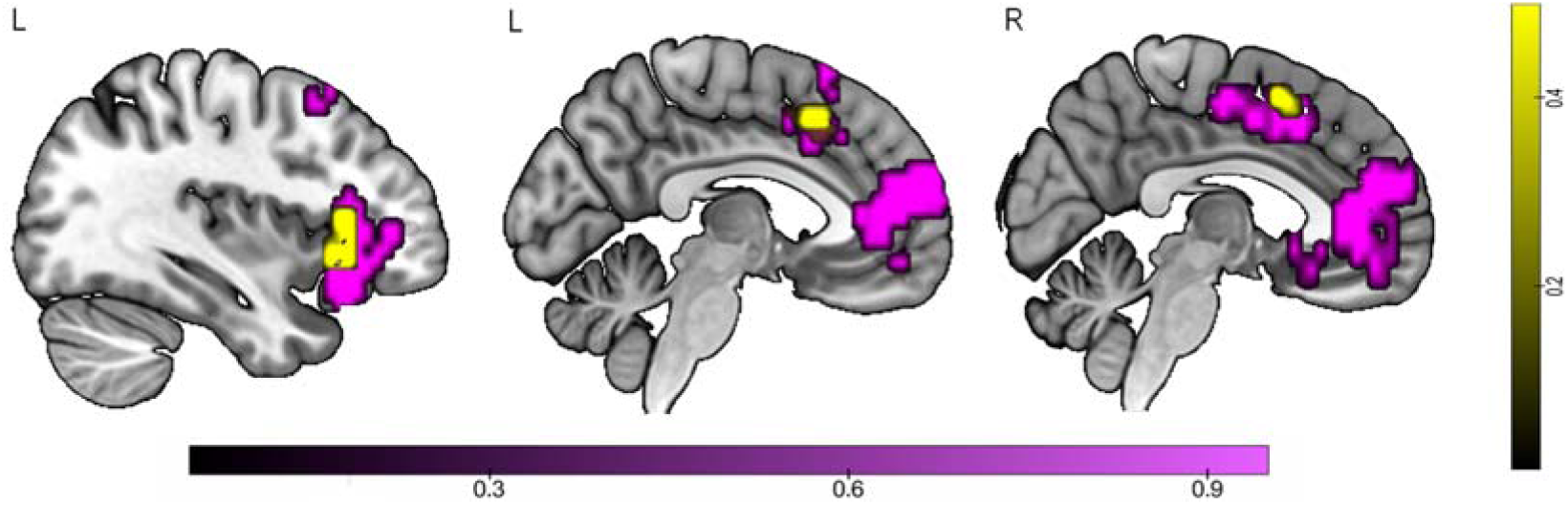
Multivariate results (violet). Different neural patterns for the valence of the adjective (positive vs. negative) during the expectation stage. Scales reflect corrected p-values (<.05). Significant regions in both univariate and multivariate analyses are highlighted in yellow.

Although the same comparisons (positive vs. negative) in univariate GLM only yielded a significant cluster activation in the IPL for positive > negative expectations, we ran a conjunction analysis (Nichols, Brett, Andersson, Wager & Poline, 2005) to test whether the regions that increased their activation during the presentation of the adjectives (positive & negative > neutral) were similar to those that contained relevant information about the valence (as reflected by multivariate results). For this, we computed the intersection between the group maps from both contrasts. Results showed two clusters (see Figure 5): one in the left IFG/aI (*k* = 56; -36, 25, 0) and one involving the bilateral SMA (*k* = 69; -8, 18, 46).

Moreover, the valence of the partners’ descriptions influenced participants’ choices, where they accepted more offers after positive than negative descriptions. As explained in the methods section (2.6 Multivariate analyses), information about participants’ responses might be employed to decode the valence of partners’ descriptions. To examine whether the regions containing relevant information about the valence were the same as those representing the participants’ decisions (accept vs. reject), we performed a response classification at the offer period and ran a conjunction analysis. Here, we observed that only a cluster in the bilateral SMA (*k* = 95; -1, 7, 48) resulted significant for both classification analyses. Additionally, we carried out a cross-classification analysis (Kaplan et al., 2015) to examine the overlap between the neural representations of participants’ choices and the valence of partners’ descriptions. In this case, that a classifier trained with response data is not able to decode valence category accurately would suggest that the neural codes underlying valence and response classifications are different and, therefore, that the valence decoding results are not explained by participants’ responses. Results from this analysis showed that cross-decoding was only possible from bilateral SMA extending to left parietal lobe (*k* = 671; -1, -11, 45), as well as from a cluster in left cerebellum extending to lingual and fusiform gyri (*k* = 381; -18, -60, -15). This indicates that classification of valence in IFG/aI and vmPFC/ACC cannot be explained by the patterns related to participants’ responses.

##### Correlation between decoding accuracy and the bias index

To explore how much influence the valence of the adjectives had on choices, we correlated the mean decoding accuracies (positive vs. negative) for each significant cluster in the MVPA with the behavioural bias index for each participant. This analysis yielded significant positive correlations between the decoding accuracy for the descriptions’ valence and the behavioural bias in all 3 significant clusters (see Figure 6): the left IFG/MFG and aI (*r* = .42; *p* = .02), bilateral vmPFC/ACC (*r* = .44; *p* = .015), and the left MCC/SMA (*r* = .53; *p* = .0038). Hence, the better the activation patterns in these regions discriminated between the valence of the partners’ information, the larger the effect of valenced information on subsequent behavioural choices. A second correlation control analysis showed that this link was not contaminated by participants’ motor responses, since there was no correlation between any of the ROIs mean accuracies and general acceptance rate per participant (all *p*s>.39), which supports the specificity of the link between valenced expectations and choices.

**Figure 6.**
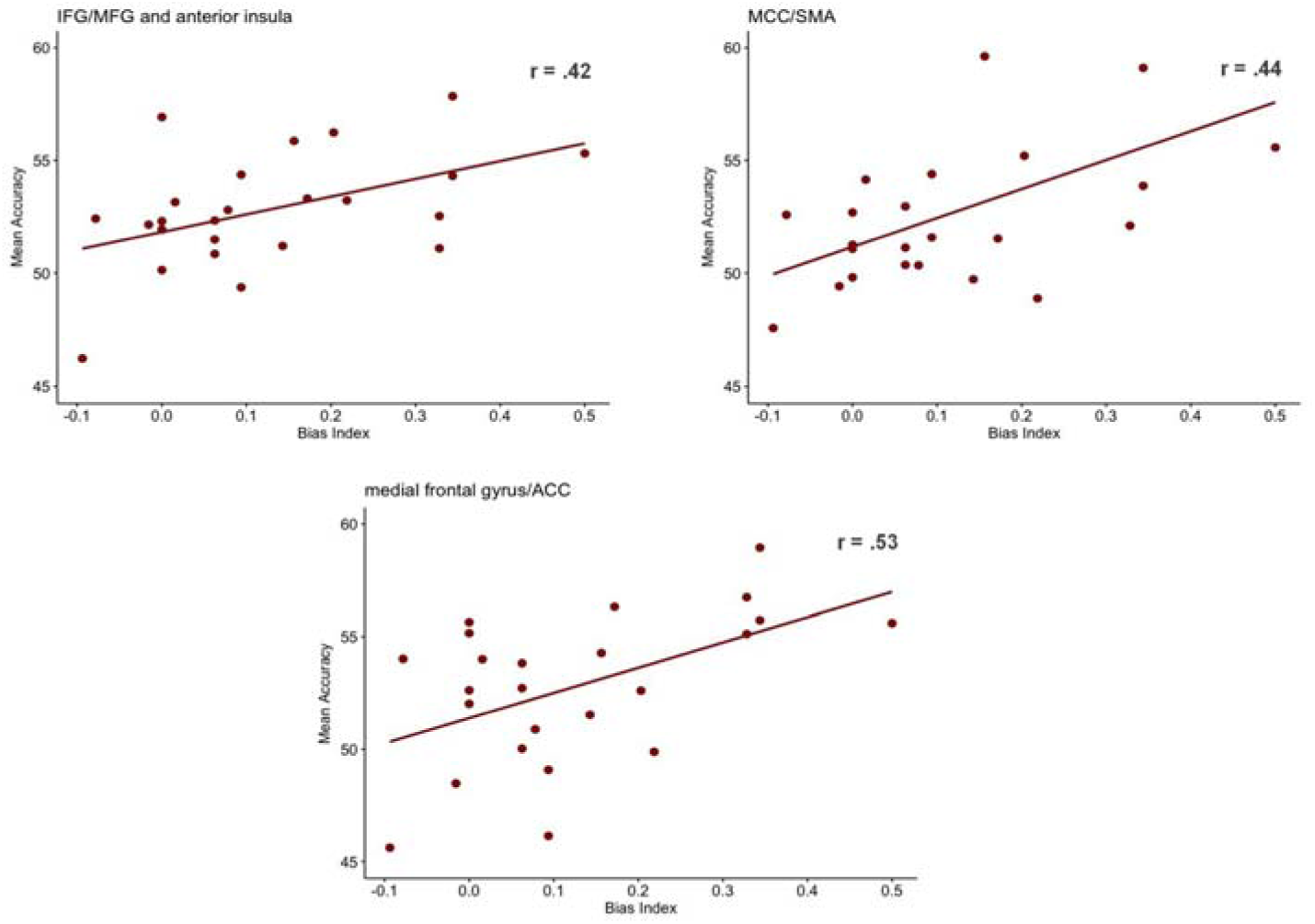
Scatter plots showing significant correlations between mean decoding accuracies in each cluster and the behavioural index. IFG: Inferior frontal gyrus. MFG: Middle frontal gyrus. ACC: Anterior Cingulate Cortex. MCC: Middle Cingulate Cortex. SMA: Supplementary Motor Area.

## 4. Discussion

Our study investigated the neural basis of social valenced expectations during an interpersonal UG. Results revealed that social information about other people bias subsequent economic choices, as well as it increases activity in the anterior insula and SMA. Furthermore, decoding analysis allowed to observe that these areas, together with the vmPFC, represent the content of such expectations. Notably, the better this information is represented in these regions, the more biased are participants to employ such knowledge when making their economic decisions.

The UG employed showed a clear behavioural effect of interpersonal expectations, where positive descriptions of others led to higher acceptance rates compared to negative ones. Additionally, the impact of the expectations was reflected on the speed of choices, where people needed more time to reject offers after positive (or neutral) expectations. This pattern indicates that participants integrate social information in their decision-making process, showing a tendency to process offers as fairer when the partner is described positively. Further, this data replicates previous results (Gaertig et al., 2012; Moser et al., 2014; Ruz et al., 2011), emphasizing the role of expectations (Sanfey, 2009) and valenced morality in decision-making (Barrett & Bliss-Moreau, 2009). Overall, the behavioural pattern of choices observed supports the utility of the experimental paradigm to induce interpersonal valenced expectations about others that bias subsequent choices made to the same set of objective behaviour (offers made by partners).

Several regions increased their activation when participants held in mind social expectations about game partners. This information engaged the SMA and the dorsal aI, which were also active at the offer stage. These are regions have been previously related to preparation processes (Brass & von Cramon, 2004), as well as sustained (Dosenbach, Fair, Cohen, Schlaggar, & Petersen, 2008; Palenciano, González-García, Arco, & Ruz, 2019) and transient (Menon & Uddin, 2010; Sridharan, Levitin, & Menon, 2008) top-down control, in paradigms where participants use cue-related information to perform tasks of different nature on subsequent targets. In previous studies using the UG, these regions have been linked to response to unfairness (Gabay et al., 2014). In addition, previous work has related aI activation with the rejection of unfair offers (Sanfey et al., 2003). In the current context, these areas may be involved in using the interpersonal information contained in the cue to guide or bias the action towards a certain choice, according to the valence of the expectation. However, univariate contrasts between the words containing positive vs. negative information, in stark contrast with behavioural outcomes, showed effects restricted on a cluster in the IPL. This region has been related to the simulation of others’ action in shared representations (Van Overwalle, 2009), and a part of our cluster it is included in the TPJ (e.g., Scholz et al., 2009), which plays a main role in ToM (Saxe & Kanwisher, 2003). The increase of activation in this region for positive expectations could indicate a higher reliance on positive descriptions by the ToM processes involved in our task. This fits with the pattern found in RTs where only positive expectations speeded acceptance choices, whereas negative descriptions did not speed rejections. Further research will be needed to replicate this imbalance of information and to better understand the nature of the underlying brain processes.

Importantly, the use of a multivariate classification analysis (MVPA) unveiled the brain regions that contain differential patterns for positive vs. negative expectations about partners. This is especially relevant since previous work has indicated how valence differences at a neural level are particularly hard to observe (Lindquist et al., 2015). These areas included the SMA/MCC, IFG/MPFC and vmPFC/ACC. There was no difference in RT between positive and negative conditions (see Behavioural data, section 3.1.), which rules out the possibility that the classifier was mistakenly discriminating faster vs. slower conditions.

The relevance of the SMA in social scenarios has been reported previously (Chang & Sanfey, 2013). These authors observed a relationship between the activity in this area and the deviation of previous expectations. Moreover, Lindquist et al. (2015) linked this region to the unspecific representation of valence. Our conjunction analysis shows that part of the SMA increases its activity during the expectation period and also shows different patterns depending on the valence of the expectation. This data suggests that the SMA has a role in general preparation but it also contains specific fine information relevant to the task. In addition, we observe partial overlapping activation with the response classification, which suggests that this region also contains some information about participants’ responses. The MCC, on the other hand, has been associated with an increase of the efficiency in decision-making, being involved in the anticipation and consequent expectations of outcomes in a variety of non-social tasks (Vogt, 2016). Further, it has also been related to the prediction and monitoring of outcomes in social decisions (Apps, Lockwood, & Balsters, 2013), and it may play a similar role in our study.

On the other hand, the patterns of activity in a lateral prefrontal cortex cluster (lPFC), including the IFG and MPFC, also discriminated the valence of the expectations. Interestingly, these areas were part of a large cluster that also increased their activation during the maintenance of social information, as revealed by univariate results. In non-social paradigms, the lPFC has been related to working memory maintenance (Morgan, Jackson, Van Koningsbruggen, Shapiro, & Linden, 2013; Sala, Rämä, & Courtney, 2003) and other forms of cognitive control (e.g., Reverberi et al., 2012). The IFG specifically has also been associated with the selection of semantic information (Jefferies, 2013; Wagner, Paré-Blagoev, Clark, & Poldrack, 2001), and it is also involved in the expectation to perform different non-social tasks employing verbal material (e.g., González-García et al., 2017; Sakai and Passingham, 2006). Notably, our results extend this role to a social context (see also Filkowski et al., 2016; Thye et al., 2018; Van Overwalle, 2009), where verbal information is used to generate positive or negative expectations about game partners, by showing that the pattern of activity in this frontal region differs depending on the nature of the information used to predict the proximal behaviour of others.

On the other hand, the vmPFC/ACC did not increase its overall activation during the expectation period but contained patterns related to the valence of the predictions. Crucially, this area overlaps with the region isolated in the meta-analysis by Lindquist et al., (2015), where they linked its activity with a bipolar representation of valence. On a broader context, this region is part of the social cognition network, associated with mentalizing processes (Koster-Hale & Saxe, 2013; Tamir et al., 2016), and behaviour guided by social cues, along with the ACC. Previous studies relate the mPFC with predictions about others’ desires (Corradi-Dell’Acqua et al., 2015), and priors during valued decisions (Lopez-Persem et al., 2016). Additionally, Van Overwalle (2009) linked this region to the integration of personal traits, and it has been extensively associated with the representation of intentions as well (Haynes et al., 2007).

The association between a brain region and a given behaviour is strengthened when a link can be observed between the fidelity of a pattern of activity and the behavioural outcome studied (Naselaris, Kay, Nishimoto, & Gallant, 2011; Tong & Pratte, 2012). To find this evidence we obtained, for each participant, a bias index representing how much the valence of the personal information influenced their choices and correlated this index with the accuracy of the classifier in disentangling the patterns generated by positive vs. negative words. We observed a positive correlation between these two factors in the three clusters sensitive to the valence of expectations. Thus, the better the classifier distinguished between descriptions of different valence, the more people tended to accept offers preceded by positive compared to negative descriptions. These results strongly suggest that these valenced representations were used to weight the posterior acceptance or rejection decisions to the same set of objective offers, biasing behaviour. Importantly, additional control correlation analysis evidenced that this finding was not contaminated by participants’ responses.

We could also observe the effect of expectations by studying the brain activity generated by offers that matched or mismatched them, that is, fair and unfair offers preceded by descriptions of the same or opposing valence. Here we found cerebellum activity when fair offers were preceded by positive descriptions and unfair ones followed negative adjectives. This region is associated with prediction in a variety of contexts, such as language (Lesage, Hansen, & Miall, 2017; Pleger & Timmann, 2018) and also social cognition (Van Overwalle, Baetens, Mariën, & Vandekerckhove, 2014), among others. In social scenarios, where people frequently anticipate others’ needs or actions, the understanding of the role of the cerebellum in predictions is particularly relevant (Sokolov, Miall, & Ivry, 2017). Although previous studies (Berthoz, 2002) found increased activity in the cerebellum when predictions (social norms) were violated, we observed the opposite. Hence, our data suggest that in the current context the cerebellum may signal when predictions are matched by social observations. Conversely, when predictions are not met, we observed activation in the lPFC, specifically the IFG and aI. In this contexts, the lFG has been associated with semantic cueing (González-García et al., 2016), semantic control (Jefferies, 2013) and emotional regulation during social decisions (Grecucci, Giorgetta, Bonini, & Sanfey, 2013). Conversely, the aI has been linked to responses to unfair offers, which represent a violation of social norms (Corradi-Dell’Acqua et al., 2013). This agrees with the incompatibility we observe here between previous expectations and actual events. Altogether, this data also supports the relevance of expectations when participants face the outcome of an interaction. At this point, they may need to suppress the previous information to act in accordance with the offer.

Although it was not the main goal of this work, we also examined brain responses to the fairness of the offer. While previous work has shown activation in areas such as aI, cingulate cortex and mPFC in reaction to unfair offers (Corradi-Dell’Acqua et al., 2013; Gabay et al., 2014), we observed higher activation in ACC/mPFC when participants faced fair (vs. unfair) offers. In this line, the mPFC has been linked to the monitoring of emotional reactions in bargaining scenarios (Corradi-Dell’Acqua et al., 2013), and its involvement could represent the positive outcome related to fair offers, in line with previous work associating the mPFC with value assessment of outcomes (Amodio & Frith, 2006). The ACC, on the other hand, has been related to the proposal of fair offers due to strategic motives (Chen, Chen, Kuo, Kan, & Yang, 2017), suggesting a role of this area in computing reward. This, in turn, would be in line with our results of the fairness of the offer, where the ACC could be relevant to signal their rewarding outcomes.

Our study has certain limitations, which should be addressed in future investigations. First, the optimal procedure to perform multivariate analyses and avoid response-related confounds is to counterbalance response options for each participant (Todd, Nystrom, & Cohen, 2013). In the current experiment, however, the association between hand and response was counterbalanced at the group but not the individual level. Thus, our valence-related classifications could have been affected by the response patterns linked to acceptance and rejection choices. To rule this out, we performed an additional conjunction analysis, which showed that only a small portion of the SMA cluster was common to both contrasts. Also, we observed that patterns in part of this region overlapped between participants’ decisions and the valence of their expectations. These results suggest that the SMA represents both events with similar codes, although it could also be the case that findings in this region are due to confounds from participants’ responses. In further support of the relevance of the representation of the valence in the bias observed in decisions, an additional control analysis showed that the performance of the classifier for the valence decoding was only related to a specific behavioural bias resulting from the valence of the expectation, but not with the response itself. Therefore, our data highlight that the fidelity of the valence representation in IFG/aI and vmPFC is associated with the extent to which the partners’ descriptions modulate participants’ decisions.

Further, it may be argued that the influence of partners’ moral information could be due to alterations in participants’ mood after reading these descriptions, rather than because the generation of expectations about their likely behaviour. Although we cannot deny completely this possibility, our findings show a specific link between participants’ behavioural bias and the neural representation of partners’ social information, which would not be in line with an explanation related to general mood fluctuations. Alternatively, following previous work on affective priming and conflict (Dignath, Eder, Steinhauser, & Kiesel, 2020; Fritz & Dreisbach, 2013), adjectives could act as affective primes (Bush et al., 2018). Although we cannot completely rule out this possibility, previous results suggest otherwise. Gaertig et al. (2012) carried out an experiment without the social cover story to test this alternative explanation. Here, the same words failed to trigger valence bias in choices. This indicates that, rather than an automatic priming effect triggered by the adjectives, it is the association between these and the character of the partners which impacted participants’ decisions. An additional concern relates to the ecological validity of our study, which is limited by the context of fMRI scanning in a single location. However, we increased the credibility of the social scenario by means of instructions and a cover story, where we recreated an actual delayed interaction between participants of different studies, and where actual earnings were contingent on the choices made during the game. In fact, none of the participants showed signs of susceptibility about the underlying nature of the study when informally debriefed at the end of the session. Nonetheless, participants could have approached the task in various ways, engaging in the social context differently. Thus, we believe that including a more detailed and structured debriefing where this and other points are addressed should be included in future studies. Moreover, another step forward would be to assess participants’ personality and prosocial tendencies, since individual predispositions can also influence these dynamics (Díaz-Gutiérrez et al., 2017). Futures studies could use some form of virtual reality during scanning (Mueller et al., 2012) together with more complex verbal descriptions of others to examine whether similar brain regions represent this content and the way this is structured, perhaps employing neuroimaging methods with higher ecological validity (e.g. Pinti et al., 2018). Additionally, another interesting research question would be to find if there is a sort of “common valence space” for the two stages of the paradigm. That is, to find out if there is shared information underlying the valence of the adjective (positive/negative) but also the “pleasantness” of the offer (fair-positive, unfair-negative). A future study designed to employ cross-classification decoding approaches (Kaplan et al., 2015) between the expectation and the evidence game periods with temporally precise methods such as electroencephalography could offer valuable information on this respect.

## Acknowledgments

This work was supported through grants by the Spanish Ministry of Science and Innovation (PSI2013-45567-P and PSI2016-78236-P to M.R.), the Spanish Ministry of Education, Culture and Sports (FPU2014/04272 to P.D.G.) and the University of Granada, through a “Contratos puente” scholarship to P.D.G.

## Notes

### Competing Interest Statement

The authors have declared no competing interest.

## References

Abdulrahman, H., & Henson, R. N. (2016). Effect of trial-to-trial variability on optimal event-related fMRI design: Implications for Beta-series correlation and multi-voxel pattern analysis. NeuroImage, 125, 756–766. https://doi.org/10.1016/j.neuroimage.2015.11.009

Amodio, D. M., & Frith, C. D. (2006). Meeting of minds: The medial frontal cortex and social cognition. Nature Reviews Neuroscience, 7(4), 268–277. https://doi.org/10.1038/nrn1884

Apps, M. A. J., Lockwood, P. L., & Balsters, J. H. (2013). The role of the midcingulate cortex in monitoring others’ decisions. Frontiers in Neuroscience, 7, 251. https://doi.org/10.3389/fnins.2013.00251

Arco, J. E., González-García, C., Díaz-Gutiérrez, P., Ramírez, J., & Ruz, M. (2018). Influence of activation pattern estimates and statistical significance tests in fMRI decoding analysis.

Barrett, L. F., & Bliss-Moreau, E. (2009). Affect as a psychological primative. Advances in Experimental Social Psychology, 41(08), 167–218. https://doi.org/10.1016/S0065-2601(08)00404-8.Affect

Berthoz, S. (2002). An fMRI study of intentional and unintentional (embarrassing) violations of social norms. Brain, 125(8), 1696–1708. https://doi.org/10.1093/brain/awf190

Bode, S., & Haynes, J. D. (2009). Decoding sequential stages of task preparation in the human brain. NeuroImage, 45(2), 606–613. https://doi.org/10.1016/j.neuroimage.2008.11.031

Brañas-Garza, P., Espín, A. M., Exadaktylos, F., & Herrmann, B. (2014). Fair and unfair punishers coexist in the Ultimatum Game. Scientific Reports, 4, 6025. https://doi.org/10.1038/srep06025

Brass, M., & von Cramon, D. Y. (2004). Decomposing Components of Task Preparation with Functional Magnetic Resonance Imaging. Journal of Cognitive Neuroscience, 16(4), 609–620. https://doi.org/10.1162/089892904323057335

Bush, K. A., Gardner, J., Privratsky, A., Chung, M. H., James, G. A., & Kilts, C. D. (2018). Brain states that encode perceived emotion are reproducible but their classification accuracy is stimulus-dependent. Frontiers in Human Neuroscience, 12(July), 1–15. https://doi.org/10.3389/fnhum.2018.00262

Chang, L. J., & Sanfey, A. G. (2013). Great expectations: Neural computations underlying the use of social norms in decision-making. Social Cognitive and Affective Neuroscience, 8(3), 277–284. https://doi.org/10.1093/scan/nsr094

Chen, Y., Chen, Y., Kuo, W., Kan, K., & Yang, C. C. (2017). Strategic Motives Drive Proposers to Offer Fairly in Ultimatum Games: An fMRI Study. Scientific Reports, 7(527), 1–11. https://doi.org/10.1038/s41598-017-00608-8

Contreras, J. M., Banaji, M. R., & Mitchell, J. P. (2012). Dissociable neural correlates of stereotypes and other forms of semantic knowledge. Social Cognitive and Affective Neuroscience, 7(7), 764–770. https://doi.org/10.1093/scan/nsr053

Contreras, J. M., Banaji, M. R., & Mitchell, J. P. (2013). Multivoxel Patterns in Fusiform Face Area Differentiate Faces by Sex and Race. PLoS ONE, 8(7), e69684. https://doi.org/10.1371/journal.pone.0069684

Corradi-Dell’Acqua, C., Civai, C., Rumiati, R. I., & Fink, G. R. (2013). Disentangling self- and fairness-related neural mechanisms involved in the ultimatum game: An fMRI study. Social Cognitive and Affective Neuroscience, 8(4), 424–431. https://doi.org/10.1093/scan/nss014

Corradi-Dell’Acqua, C., Turri, F., Kaufmann, L., Clément, F., & Schwartz, S. (2015). How the brain predicts people’s behavior in relation to rules and desires. Evidence of a medio-prefrontal dissociation. Cortex, 70, 21–34. https://doi.org/10.1016/j.cortex.2015.02.011

Correa, A., Alguacil, S., Ciria, L. F., Jiménez, A., & Ruz, M. (2020). Circadian rhythms and decision-makingLJ: a review and new evidence from electroencephalography. Chronobiology International. https://doi.org/10.1080/07420528.2020.1715421

Correa, A., Ruiz-Herrera, N., Ruz, M., Tonetti, L., Martoni, M., Fabbri, M., & Natale, V. (2017). Economic decision-making in morning/evening-type people as a function of time of day. Chronobiology International, 34(2), 139–147. https://doi.org/10.1080/07420528.2016.1246455

Delgado, M. R., Frank, R. H., & Phelps, E. A. (2005). Perceptions of moral character modulate the neural systems of reward during the trust game. Nature Neuroscience, 8(11), 1611–1618. https://doi.org/10.1038/nn1575

Di Russo, F., Berchicci, M., Bozzacchi, C., Perri, R. L., Pitzalis, S., & Spinelli, D. (2017). Beyond the “Bereitschaftspotential”: Action preparation behind cognitive functions. Neuroscience and Biobehavioral Reviews, 78(April), 57–81. https://doi.org/10.1016/j.neubiorev.2017.04.019

Díaz-Gutiérrez, P., Alguacil, S., & Ruz, M. (2017). Bias and control in social decision-making. In A. Ibáñez, L. Sedeño, & A. Gacría (Eds.), Neuroscience and Social Science: The Missing Link (pp. 47–68). Cham, Switzerland: Springer. https://doi.org/10.1007/978-3-319-68421-5

Dignath, D., Eder, A. B., Steinhauser, M., & Kiesel, A. (2020). Conflict monitoring and the affective-signaling hypothesis—An integrative review. Psychonomic Bulletin and Review, 27(2), 193–216. https://doi.org/10.3758/s13423-019-01668-9

Dosenbach, N. U. F., Fair, D. A., Cohen, A. L., Schlaggar, B. L., & Petersen, S. E. (2008). A dual-networks architecture of top-down control. Trends in Cognitive Sciences, 12(3), 99–105. https://doi.org/10.1016/j.tics.2008.01.001

Esterman, M., & Yantis, S. (2010). Perceptual expectation evokes category-selective cortical activity. Cerebral Cortex, 20(5), 1245–1253. https://doi.org/10.1093/cercor/bhp188

Fehr, E., & Camerer, C. F. (2007). Social neuroeconomics: the neural circuitry of social preferences. Trends in Cognitive Sciences, 11(10), 419–427. https://doi.org/10.1016/j.tics.2007.09.002

Filkowski, M. M., Anderson, I. W., & Haas, B. W. (2016). Trying to trust: Brain activity during interpersonal social attitude change. Cognitive, Affective and Behavioral Neuroscience, 16(2), 325–338. https://doi.org/10.3758/s13415-015-0393-0

Fleming, S. M., Thomas, C. L., & Dolan, R. J. (2010). Overcoming status quo bias in the human brain. Proceedings of the National Academy of Sciences, 107(13), 6005–6009. https://doi.org/10.1073/pnas.0910380107

Fouragnan, E., Chierchia, G., Greiner, S., Neveu, R., Avesani, P., & Coricelli, G. (2013). Reputational Priors Magnify Striatal Responses to Violations of Trust. The Journal of Neuroscience, 33(8), 3602–3611. https://doi.org/10.1523/JNEUROSCI.3086-12.2013

Friston, K. (2005). A theory of cortical responses. Philosophical Transactions of the Royal Society B: Biological Sciences, 360(1456), 815–836. https://doi.org/10.1098/rstb.2005.1622

Frith, C. D. (2007). The social brain? Phil. Trans. R. Soc. B, 362(10), 671–678. https://doi.org/10.1098/rstb.2006.2003

Frith, C. D., & Frith, U. (2008). Implicit and Explicit Processes in Social Cognition. Neuron, 60(3), 503–510. https://doi.org/10.1016/j.neuron.2008.10.032

Frith, U., & Frith, C. (2001). The biological basis of social interaction. American Psychological Society, 10(5), 151–155. https://doi.org/https://doi.org/10.1111/1467-8721.00137

Fritz, J., & Dreisbach, G. (2013). Conflicts as aversive signals: Conflict priming increases negative judgments for neutral stimuli. Cognitive, Affective and Behavioral Neuroscience, 13(2), 311–317. https://doi.org/10.3758/s13415-012-0147-1

Gabay, A. S., Radua, J., Kempton, M. J., & Mehta, M. A. (2014). The Ultimatum Game and the brain: A meta-analysis of neuroimaging studies. Neuroscience and Biobehavioral Reviews, 47, 549–558. https://doi.org/10.1016/j.neubiorev.2014.10.014

Gaertig, C., Moser, A., Alguacil, S., & Ruz, M. (2012). Social information and economic decision-making in the ultimatum game. Frontiers in Neuroscience, 6:103. https://doi.org/10.3389/fnins.2012.00103

González-García, C., Arco, J. E., Palenciano, A. F., Ramírez, J., & Ruz, M. (2017). Encoding, preparation and implementation of novel complex verbal instructions. NeuroImage, 148(January), 264–273. https://doi.org/10.1016/j.neuroimage.2017.01.037

González-García, C., Mas-Herrero, E., de Diego-Balaguer, R., & Ruz, M. (2016). Task-specific preparatory neural activations in low-interference contexts. Brain Structure and Function, 221(8), 3997–4006. https://doi.org/10.1007/s00429-015-1141-5

Grecucci, A., Giorgetta, C., Bonini, N., & Sanfey, A. G. (2013). Reappraising social emotions: the role of inferior frontal gyrus, temporo-parietal junction and insula in interpersonal emotion regulation. Frontiers in Human Neuroscience, 7:523. https://doi.org/10.3389/fnhum.2013.00523

Güth, W., Schmittberger, R., & Schwarze, B. (1982). An experimental analysis of ultimatum bargaining. Journal of Economic Behavior and Organization, 3(4), 367–388. https://doi.org/10.1016/0167-2681(82)90011-7

Hajcak, G., Moser, J. S., Holroyd, C. B., & Simons, R. F. (2006). The feedback-related negativity reflects the binary evaluation of good versus bad outcomes. Biological Psychology, 71(2), 148–154. https://doi.org/10.1016/j.biopsycho.2005.04.001

Hassabis, D., Spreng, R. N., Rusu, A. A., Robbins, C. A., Mar, R. A., & Schacter, D. L. (2014). Imagine all the people: How the brain creates and uses personality models to predict behavior. Cerebral Cortex, 24(8), 1979–1987. https://doi.org/10.1093/cercor/bht042

Haynes, J., Sakai, K., Rees, G., Gilbert, S., Frith, C., & Passingham, R. E. (2007). Reading Hidden Intentions in the Human Brain. Current Biology, 17(4), 323–328. https://doi.org/10.1016/j.cub.2006.11.072

Hebart, M. N., Görgen, K., & Haynes, J.-D. (2015). The Decoding Toolbox (TDT): a versatile software package for multivariate analyses of functional imaging data. Frontiers in Neuroinformatics, 8, 88. https://doi.org/10.3389/fninf.2014.00088

Jefferies, E. (2013). The neural basis of semantic cognition: Converging evidence from neuropsychology, neuroimaging and TMS. Cortex, 49(3), 611–625. https://doi.org/10.1016/j.cortex.2012.10.008

Kahneman, D., Knetsch, J. L., & Thaler, R. H. (1986). Fairness and Assumptions of Economics. Journal of Business, 59(4), S285–300.

Kaplan, J. T., Man, K., & Greening, S. G. (2015). Multivariate cross-classification: applying machine learning techniques to characterize abstraction in neural representations. Frontiers in Human Neuroscience, 9, 151. https://doi.org/10.3389/fnhum.2015.00151

Knoch, D., Pascual-Leone, A., Meyer, K., Treyer, V., & Fehr, E. (2006). Diminishing reciprocal fairness by disrupting the right prefrontal cortex. Science, 314(5800), 829–832. https://doi.org/10.1126/science.1129156

Koster-Hale, J., & Saxe, R. (2013). Theory of Mind: A Neural Prediction Problem. Neuron, 79(5), 836–848. https://doi.org/10.1016/j.neuron.2013.08.020

Kriegeskorte, N., Goebel, R., & Bandettini, P. (2006). Information-based functional brain mapping. Proceedings of the National Academy of Sciences of the United States of America, 103, 3863–3868. https://doi.org/10.1073/pnas.0600244103

Lesage, E., Hansen, P. C., & Miall, R. C. (2017). Right Lateral Cerebellum Represents Linguistic Predictability. The Journal of Neuroscience, 37(26), 6231–6241. https://doi.org/10.1523/JNEUROSCI.3203-16.2017

Lindquist, K. A., Satpute, A. B., Wager, T. D., Weber, J., & Barrett, L. F. (2015). The Brain Basis of Positive and Negative Affect: Evidence from a Meta-Analysis of the Human Neuroimaging Literature. Cerebral Cortex, 26(5), 1910–1922. https://doi.org/10.1093/cercor/bhv001

Lopez-Persem, A., Domenech, P., & Pessiglione, M. (2016). How prior preferences determine decision-making frames and biases in the human brain. ELife, 5: e20317. https://doi.org/10.7554/eLife.20317

Ma, N., Vandekerckhove, M., Baetens, K., Overwalle, F. Van, Seurinck, R., & Fias, W. (2012). Inconsistencies in spontaneous and intentional trait inferences. Social Cognitive and Affective Neuroscience, 7(8), 937–950. https://doi.org/10.1093/scan/nsr064

Menon, V., & Uddin, L. Q. (2010). Saliency, switching, attention and control: a network model of insula function. Brain Structure and Function, 214(5–6), 655–667. https://doi.org/10.1007/s00429-010-0262-0

Mitchell, T. M., Shinkareva, S. V., Carlson, A., Chang, K.-M., Malave, V. L., Mason, R. A., & Just, M. A. (2008). Predicting Human Brain Activity Associated with the Meanings of Nouns. Science, 320, 1191–1195. https://doi.org/10.1126/science.1152876

Morgan, H. M., Jackson, M. C., Van Koningsbruggen, M. G., Shapiro, K. L., & Linden, D. E. J. (2013). Frontal and parietal theta burst TMS impairs working memory for visual-spatial conjunctions. Brain Stimulation, 6(2), 122–129. https://doi.org/10.1016/j.brs.2012.03.001

Moser, A., Gaertig, C., & Ruz, M. (2014). Social information and personal interests modulate neural activity during economic decision-making. Frontiers in Human Neuroscience, 8: 31. https://doi.org/10.3389/fnhum.2014.00031

Mueller, C., Luehrs, M., Baecke, S., Adolf, D., Luetzkendorf, R., Luchtmann, M., & Bernarding, J. (2012). Building virtual reality fMRI paradigms: A framework for presenting immersive virtual environments. Journal of Neuroscience Methods, 209(2), 290–298. https://doi.org/10.1016/j.jneumeth.2012.06.025

Naselaris, T., Kay, K. N., Nishimoto, S., & Gallant, J. L. (2011). Encoding and decoding in fMRI. NeuroImage, 56(2), 400–410. https://doi.org/10.1016/j.neuroimage.2010.07.073

Nichols, T., Brett, M., Andersson, J., Wager, T., & Poline, J. B. (2005). Valid conjunction inference with the minimum statistic. NeuroImage, 25(3), 653–660. https://doi.org/10.1016/j.neuroimage.2004.12.005

Palenciano, A. F., González-García, C., Arco, J. E., & Ruz, M. (2019). Transient and Sustained Control Mechanisms Supporting Novel Instructed Behavior. Cerebral Cortex, 29(9), 3948–3960. https://doi.org/10.1093/cercor/bhy273

Pereira, F., Mitchell, T., & Botvinick, M. (2009). Machine learning classi!ers and fMRI: A tutorial overview. NeuroImage, 45(1), S199–S209. https://doi.org/10.1016/j.neuroimage.2008.11.007.Machine

Pinti, P., Tachtsidis, I., Hamilton, A., Hirsch, J., Aichelburg, C., Gilbert, S., & Burgess, P. W. (2018). The present and future use of functional near-infrared spectroscopy (fNIRS) for cognitive neuroscience. Annals of the New York Academy of Sciences, 1–25. https://doi.org/10.1111/nyas.13948

Pleger, B., & Timmann, D. (2018). The role of the human cerebellum in linguistic prediction, word generation and verbal working memory: evidence from brain imaging, non-invasive cerebellar stimulation and lesion studies. Neuropsychologia. https://doi.org/10.1016/j.neuropsychologia.2018.03.012

Puri, A. M., Wojciulik, E., & Ranganath, C. (2009). Category expectation modulates baseline and stimulus-evoked activity in human inferotemporal cortex. Brain Research, 1301, 89–99. https://doi.org/10.1016/j.brainres.2009.08.085

Redondo, J., Fraga, I., Padrón, I., & Comesaña, M. (2007). The Spanish adaptation of anew (Affective Norms for English Words). Behavior Research Methods, 39(3), 600–605. https://doi.org/10.3758/BF03193031

Reverberi, C., Görgen, K., & Haynes, J.-D. (2012). Compositionality of Rule Representations in Human Prefrontal Cortex. Cerebral Cortex, 22(6), 1237–1246. https://doi.org/10.1093/cercor/bhr200

Ruz, M., Moser, A., & Webster, K. (2011). Social expectations bias decision-making in uncertain inter-personal situations. PLoS ONE, 6(2): e157. https://doi.org/10.1371/journal.pone.0015762

Ruz, M., & Tudela, P. (2011). Emotional conflict in interpersonal interactions. NeuroImage, 54(2), 1685–1691. https://doi.org/10.1016/j.neuroimage.2010.08.039

Sakai, K., & Passingham, R. E. (2006). Prefrontal Set Activity Predicts Rule-Specific Neural Processing during Subsequent Cognitive Performance. Journal of Neuroscience, 26(4), 1211–1218. https://doi.org/10.1523/JNEUROSCI.3887-05.2006

Sakai, Katsuyuki. (2008). Task Set and Prefrontal Cortex. Annual Review of Neuroscience, 31(1), 219–245. https://doi.org/10.1146/annurev.neuro.31.060407.125642

Sala, J. B., Rämä, P., & Courtney, S. M. (2003). Functional topography of a distributed neural system for spatial and nonspatial information maintenance in working memory. Neuropsychologia, 41(3), 341–356. https://doi.org/10.1016/S0028-3932(02)00166-5

Sanfey, A. G. (2009). Expectations and social decision-making: Biasing effects of prior knowledge on Ultimatum responses. Mind and Society, 8(1), 93–107. https://doi.org/10.1007/s11299-009-0053-6

Sanfey, A. G., Rilling, J. K., Aronson, J. A., Nystrom, L. E., & Cohen, J. D. (2003). The neural basis of economic decision-making in the Ultimatum Game. Science, 300, 1755–1758. https://doi.org/10.1126/science.1082976

Saxe, R., & Kanwisher, N. (2003). People thinking about thinking peopleThe role of the temporo-parietal junction in “theory of mind.” NeuroImage, 19(4), 1835–1842. https://doi.org/10.1016/S1053-8119(03)00230-1

Schneider, W., Eschman, A., & Zuccolotto, A. (2002). E-Prime user’s guide. Pittsburgh: Psychology Software Tools Inc.

Scholz, J., Triantafyllou, C., Whitfield-Gabrieli, S., Brown, E. N., & Saxe, R. (2009). Distinct regions of right temporo-parietal junction are selective for theory of mind and exogenous attention. PLoS ONE, 4(3). https://doi.org/10.1371/journal.pone.0004869

Schwarz, K. A., Pfister, R., & Büchel, C. (2016). Rethinking Explicit Expectations: Connecting Placebos, Social Cognition, and Contextual Perception. Trends in Cognitive Sciences, 20(6), 469–480. https://doi.org/10.1016/j.tics.2016.04.001

Sokolov, A. A., Miall, R. C., & Ivry, R. B. (2017). The Cerebellum: Adaptive Prediction for Movement and Cognition. Trends in Cognitive Sciences, 21(5), 313–332. https://doi.org/10.1016/j.tics.2017.02.005

Sridharan, D., Levitin, D. J., & Menon, V. (2008). A critical role for the right fronto-insular cortex in switching between central-executive and default-mode networks. Proceedings of the National Academy of Sciences, 105(34), 12569–12574. https://doi.org/10.1073/pnas.0800005105

Stelzer, J., Chen, Y., & Turner, R. (2013). Statistical inference and multiple testing correction in classification-based multi-voxel pattern analysis (MVPA): Random permutations and cluster size control. NeuroImage, 65, 69–82. https://doi.org/10.1016/j.neuroimage.2012.09.063

Stolier, R. M., & Freeman, J. B. (2016). Neural pattern similarity reveals the inherent intersection of social categories. Nature Neuroscience, 19(6), 795–797. https://doi.org/10.1038/nn.4296

Stolier, R. M., & Freeman, J. B. (2017). A Neural Mechanism of Social Categorization. The Journal of Neuroscience, 37(23), 5711–5721. https://doi.org/10.1523/JNEUROSCI.3334-16.2017

Summerfield, C., & De Lange, F. P. (2014). Expectation in perceptual decision making: Neural and computational mechanisms. Nature Reviews Neuroscience, 15(11), 745–756. https://doi.org/10.1038/nrn3838

Tamir, D. I., & Thornton, M. A. (2018). Modeling the Predictive Social Mind. Trends in Cognitive Sciences, 22(3), 201–212. https://doi.org/10.1016/j.tics.2017.12.005

Tamir, D. I., Thornton, M. A., Contreras, J. M., & Mitchell, J. P. (2016). Neural evidence that three dimensions organize mental state representation: Rationality, social impact, and valence. Proceedings of the National Academy of Sciences of the United States of America, 113(1), 194–199. https://doi.org/10.1073/pnas.1511905112

Thornton, M. A., & Mitchell, J. P. (2017). Theories of Person Perception Predict Patterns of Neural Activity During Mentalizing. Cerebral Cortex, 1–16. https://doi.org/10.1093/cercor/bhx216

Thye, M. D., Murdaugh, D. L., & Kana, R. K. (2018). Brain Mechanisms Underlying Reading the Mind from Eyes, Voice, and Actions. Neuroscience, 374, 172–186. https://doi.org/10.1016/j.neuroscience.2018.01.045

Todd, M. T., Nystrom, L. E., & Cohen, J. D. (2013). Confounds in multivariate pattern analysis: Theory and rule representation case study. NeuroImage, 77, 157–165. https://doi.org/10.1016/j.neuroimage.2013.03.039

Tong, F., & Pratte, M. S. (2012). Decoding Patterns of Human Brain Activity. Annual Review of Psychology, 63(1), 483–509. https://doi.org/10.1146/annurev-psych-120710-100412

Turner, B. (2010). Comparison of methods for the use of pattern classificaion on rapid event-related fMRI data. Poster session presented at the Annual Meeting of the Society for Neuroscience, San Diego, CA.

Van Overwalle, F. (2009). Social cognition and the brain: A meta-analysis. Human Brain Mapping, 30(3), 829–858. https://doi.org/10.1002/hbm.20547

Van Overwalle, F., Baetens, K., Mariën, P., & Vandekerckhove, M. (2014). Social cognition and the cerebellum: A meta-analysis of over 350 fMRI studies. NeuroImage, 86, 554–572. https://doi.org/10.1016/j.neuroimage.2013.09.033

Vogt, B. A. (2016). Midcingulate cortex: Structure, connections, homologies, functions and diseases. Journal of Chemical Neuroanatomy, 74, 28–46. https://doi.org/10.1016/j.jchemneu.2016.01.010

Wagner, A. D., Paré-Blagoev, E. J., Clark, J., & Poldrack, R. A. (2001). Recovering meaning: Left prefrontal cortex guides controlled semantic retrieval. Neuron, 31(2), 329–338. https://doi.org/10.1016/S0896-6273(01)00359-2

Yeung, N., & Sanfey, A. G. (2004). Independent Coding of Reward Magnitude and Valence in the Human Brain. Journal of Neuroscience, 24(28), 6258–6264. https://doi.org/10.1523/JNEUROSCI.4537-03.2004

